# Multiple sclerosis-associated changes in the composition and immune functions of spore-forming bacteria

**DOI:** 10.1101/343558

**Authors:** Egle Cekanaviciute, Anne-Katrin Pröbstel, Anna Thomann, Tessel F. Runia, Patrizia Casaccia, Ilana Katz Sand, Elizabeth Crabtree, Sneha Singh, John Morrissey, Patrick Barba, Refujia Gomez, Rob Knight, Sarkis K. Mazmanian, Jennifer Graves, Bruce A.C. Cree, Scott S. Zamvil, Sergio E. Baranzini

## Abstract

Multiple sclerosis (MS) is an autoimmune disease of the central nervous system characterized by adaptive and innate immune system dysregulation. Recent work has revealed moderate alteration of gut microbial communities in subjects with MS and in experimental, induced models. However, a mechanistic understanding linking the observed changes in the microbiota and the presence of the disease is still missing. Chloroform-resistant, spore-forming bacteria have been shown to exhibit immunomodulatory properties in vitro and in vivo, but they have not yet been characterized in the context of human disease. This study addresses the community composition and immune function of this bacterial fraction in MS. We identify MS-associated spore-forming taxa and show that their presence correlates with impaired differentiation of IL-10 secreting, regulatory T lymphocytes *in-vitro*. Colonization of antibiotic-treated mice with spore-forming bacteria allowed us to identify some bacterial taxa favoring IL-10^+^ lymphocyte differentiation and others inducing differentiation of pro-inflammatory, IFNγ^+^ T lymphocytes. However, when fed into antibiotic-treated mice, both MS and control derived spore-forming bacteria were able to induce immunoregulatory responses.

Our analysis also identified *Akkermansia muciniphila* as a key organism that may interact either directly or indirectly with spore-forming bacteria to exacerbate the inflammatory effects of MS-associated gut microbiota. Thus, changes in the spore-forming fraction may influence T lymphocyte-mediated inflammation in MS. This experimental approach of isolating a subset of microbiota based on its functional characteristics may be useful to investigate other microbial fractions at greater depth.

**Importance:** Despite the rapid emergence of microbiome related studies in human diseases, few go beyond a simple description of relative taxa levels in a select group of patients. Our study integrates computational analysis with *in vitro* and *in vivo* exploration of inflammatory properties of both complete microbial communities and individual taxa, revealing novel functional associations. We specifically show that while small differences exist between the microbiomes of MS patients and healthy subjects, these differences are exacerbated in the chloroform resistant fraction. We further demonstrate that, when purified from MS patients, this fraction is associated with impaired immunomodulatory responses in vitro.

## Introduction

The human gut microbiota is emerging as a major immune regulator in health and disease, particularly in relation to autoimmune disorders. Most human microbiota studies to date have been based on unbiased exploration of complete microbial communities. However, limited sequencing depth, combined with high community richness and natural sample heterogeneity, might hinder the discovery of physiologically relevant taxonomical differences. Thus, targeted studies of specific microbial populations with defined characteristics may serve as a complementary approach to investigate disease-associated changes in gut microbiome.

Spore-forming bacteria constitute a subset of Gram-positive bacteria that are resistant to 3% chloroform treatment (1, 2). Both human and mouse spore-forming bacteria have immunoregulatory functions (3, 4). Mouse spore-forming bacteria include segmented filamentous bacteria and *Clostridia* species, which have been shown to induce gut T helper lymphocyte responses (3, 5). More recently, human spore-forming bacteria from a healthy subject were also reported to induce Tregs *in vitro* and in gnotobiotic mice *(4)*. However, whether the composition and functions of spore-forming bacteria are altered in immune mediated diseases is unknown.

Multiple sclerosis (MS) is a chronic disease of the central nervous system, characterized by autoimmune destruction of myelin. MS pathogenesis is in part mediated by effector T lymphocytes, and counterbalanced by Tregs, that limit the autoimmune damage inflicted by the former population (6, 7) and potentially promote remyelination (8). Recent studies, including our own, associated MS with moderate changes in the relative amounts of gut microbiota that exacerbate T lymphocyte-mediated inflammation *in vitro* and *in vivo* by stimulating pro-inflammatory IFNγ+ Th1 and inhibiting IL-10+ regulatory T lymphocytes (9, 10).

We hypothesized that these MS-associated changes in gut microbial communities may involve spore-forming bacteria thus altering its overall immunoregulatory properties. To address this hypothesis, we isolated spore-forming bacteria from untreated patients with relapsing-remitting MS (RRMS) and matched controls and analyzed their community composition and immunoregulatory functions *in vitro* and in the experimental autoimmune encephalomyelitis (EAE) mouse model.

## Results

### MS-associated differences in microbial community composition are more evident in the spore-forming fraction

We isolated the spore-forming bacterial fraction from stool samples of 25 untreated MS patients and 24 controls and tested their relative abundance by amplicon sequencing of 16S rRNA V4 gene sequencing. This analysis revealed no differences in community richness between patients and controls (Chao1 metric of alpha diversity, Fig. 1A). However, a focused analysis on the spore-forming fraction increased sample variability both within and between groups (Fig. 1B) possibly by reducing the number of taxa of interest and thus amplifying the differences in their relative abundances (Fig. 1A, 1C).

**Figure 1.**
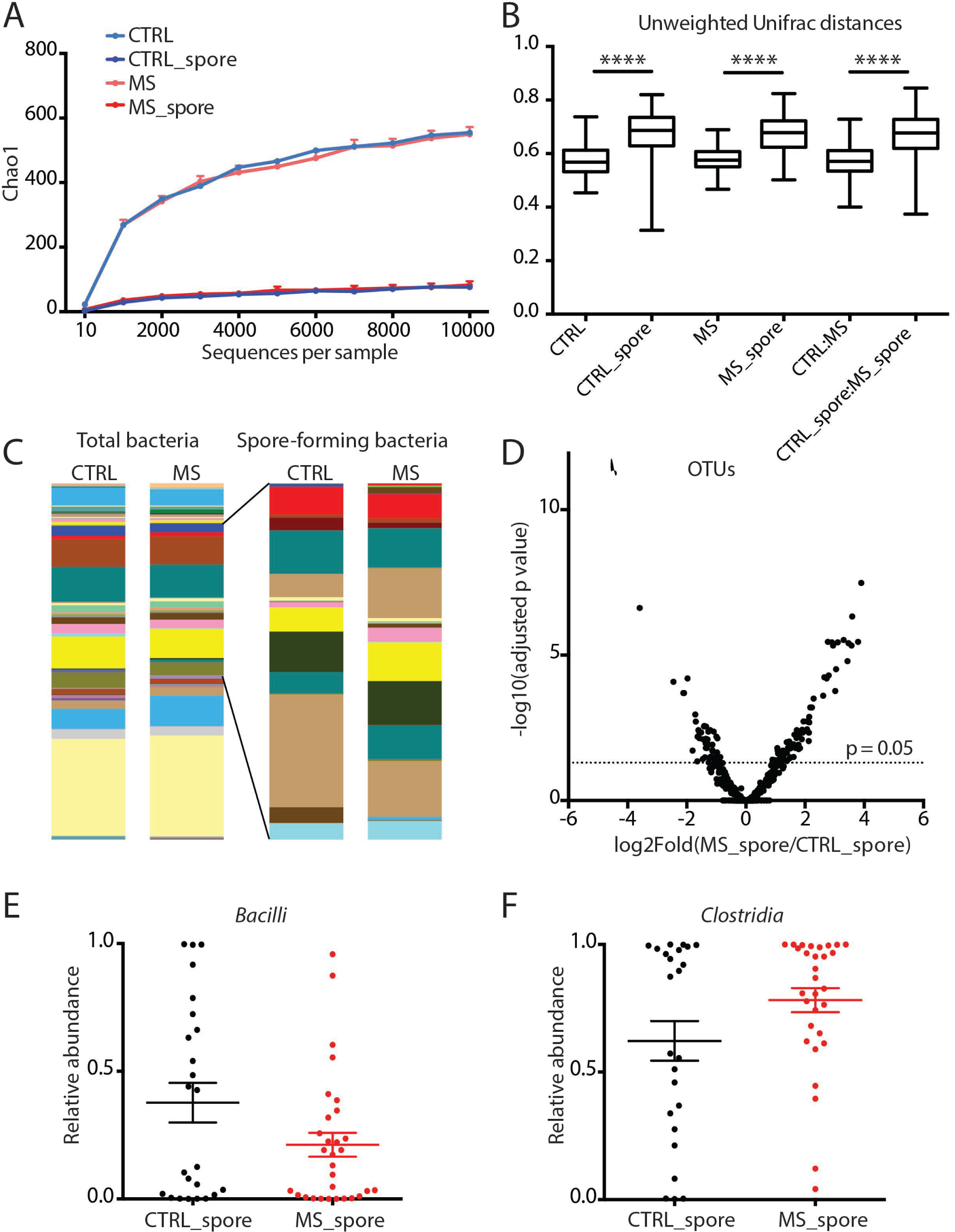
Differences in community composition of spore-forming bacterial fraction in MS patients and healthy controls. **A-C.** Comparison of microbial community composition of spore-forming bacterial subset and total stool bacteria in untreated MS patients (n=25) and controls (n=24). **A**. Chao1 metric of alpha diversity. **B.** Median and range of distances (unweighted Unifrac distance matrix) within and between sample groups. **C**. Mean relative abundance of microbial genera. **D-F.** Comparison of relative abundances of individual microbial taxa in untreated MS patients (n=25) and controls (n=24). **D.** Volcano plot of relative abundance distribution of microbial OTUs. X axis, log2 fold of relative abundance ratio between MS patients and controls after variance-stabilizing transformation. Y axis, negative log10 of P value, negative binomial Wald test, Benjamini-Hochberg correction for multiple comparisons. **E, F.** Relative abundances of bacterial classes *Bacilli* (**E**) and *Clostridia* (**F**) within phylum *Firmicutes* out of spore-forming bacteria from controls and MS patients. Error bars, mean +/- SEM. CTRL, total stool bacteria from controls. CTRL_spore, spore-forming bacteria from controls. MS, total stool bacteria from MS patients. MS_spore, spore-forming bacteria from MS patients.

Spore-forming bacteria showed notable differences in taxonomical composition between cases and controls, with 22.43% (135 out of 602 total) OTUs significantly different (p=0.05, negative binomial Wald test, Benjamini-Hochberg correction) (Fig. 1D and Suppl. Table 1). These taxonomical differences were also noticeable at the class level in which *Bacilli* were significantly overrepresented in controls (Fig. 1E), and *Clostridia*, including *Clostridium perfringens* were significantly overrepresented in MS patients (Fig. 1F and Suppl. Fig. S1).

### Spore-forming bacteria from MS patients fail to induce anti-inflammatory T lymphocytes *in vitro*

To investigate whether MS-associated differences in community composition of spore-forming bacteria were sufficient to alter the immune functions of primary blood mononuclear cells (PBMCs) from healthy human donors, we exposed human PBMCs to extracts of spore-forming bacteria isolated either from unrelated controls or from MS patients and used flow cytometry to evaluate T lymphocyte differentiation under different polarizing conditions (18-20). A comparison of the PBMC response to extracts of spore-forming bacteria from controls or from MS patients identified lower conversion into CD4+FoxP3+ Tregs (Fig. 2 A, C), including IL-10 expressing Treg population (Fig. 2 B, D) in the PBMCs exposed to the MS-derived spore-forming bacteria. These data suggest that spore-forming bacteria from MS patients are significantly less effective at inducing Treg differentiation. Of note, the small population of Tregs that still differentiated in response to MS bacteria, retained their suppressive capacities *in vitro* (Fig. 2E), thereby indicating that this was a functionally active population. Interestingly, the percentage of IL-10+ Tregs induced by extracts of spore-forming bacteria positively correlated with the relative abundance of *Bacilli* and negatively correlated with the relative abundance of *Clostridia* (Fig. 2 F, expressed as *Clostridia-Bacilli* difference). Thus, the community composition of spore-forming bacteria (i.e. high *Clostridia,* low *Bacilli*) associated with MS was also correlated with an inhibition of their respective immunoregulatory functions.

**Figure 2.**
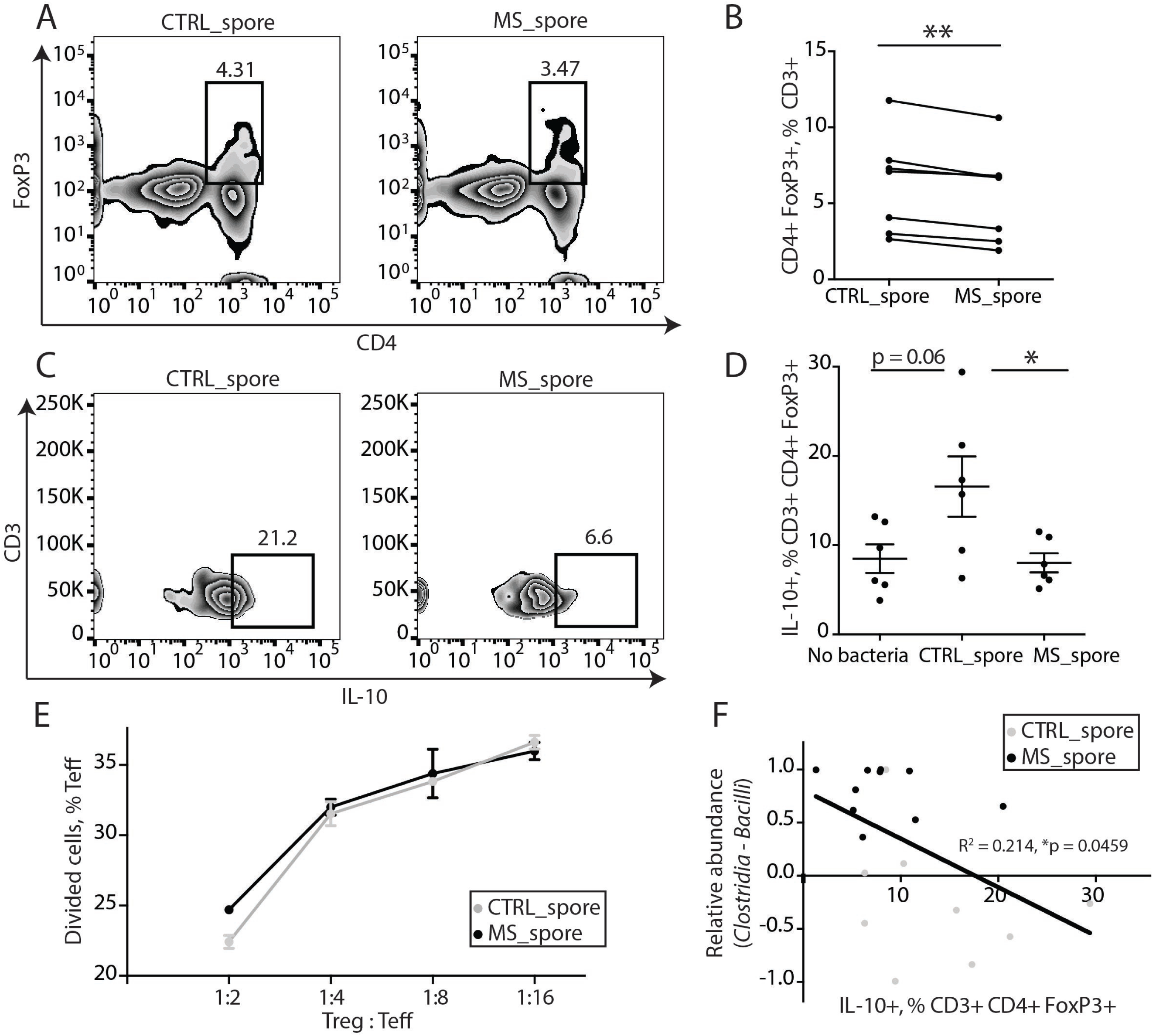
Spore-forming bacteria from MS patients inhibit IL-10+ Treg differentiation *in vitro*. **A, B.** Representative flow cytometry plots (**A**) and quantification (**B**) of CD4+FoxP3+ Tregs within CD3+ lymphocytes differentiated in response to spore-forming bacteria isolated from controls or untreated MS patients. N = 7 PBMC donors; each dot represents an average response from PBMC donor to isolates from 6 control or MS bacteria donors. **P<0.01, two-tailed repeated measures t test. **C, D.** Representative flow cytometry plots (**C**) and quantification (**D**) of IL-10+ lymphocyte population within CD3+CD4+FoxP3+ Tregs differentiated in response to spore-forming bacteria isolated from controls or untreated MS patients. N = 6 bacteria donors per group. *P<0.05, two-tailed t test. Error bars, mean +/- SEM. Experiment was repeated with non-overlapping PBMC and bacterial donors and gave the same results. **E.** Quantification of T effector cell proliferation in response to Tregs differentiated in presence of spore-forming bacteria from MS patients or controls. N = 3 bacterial donors per group, each representing an average of 3 technical replicates. **F** Linear correlation between IL-10+ population within CD3+ CD4+ FoxP3+ Tregs and *Clostridia-Bacilli* relative abundances. R^2^ = 0.214, p = 0.0459. Red dots, MS patients. Blue dots, controls. R^2 = 0.214, p = 0.046.

### Gnotobiotic mouse models reveal associations between individual bacterial taxa and T lymphocyte responses

To determine whether the MS-associated reduction in the ability of spore-forming bacteria to stimulate Treg differentiation was physiologically significant, we colonized a group of female antibiotic-treated mice (21) with spore-forming bacteria from either controls (n=2) or MS subjects (n=2) and measured the course and severity of EAE. We observed a significant reduction in disease severity in all mice whose GI tracts were reconstituted with spore-forming bacteria. However, this reduction was independent of whether the spore-forming fraction was isolated from MS or controls (Fig. 3 A). This indicated that while MS-derived spore-forming bacteria could be functionally distinguished *in vitro*, these differences were not sufficient to induce a phenotype *in vivo* in our experimental setting.

**Figure 3.**
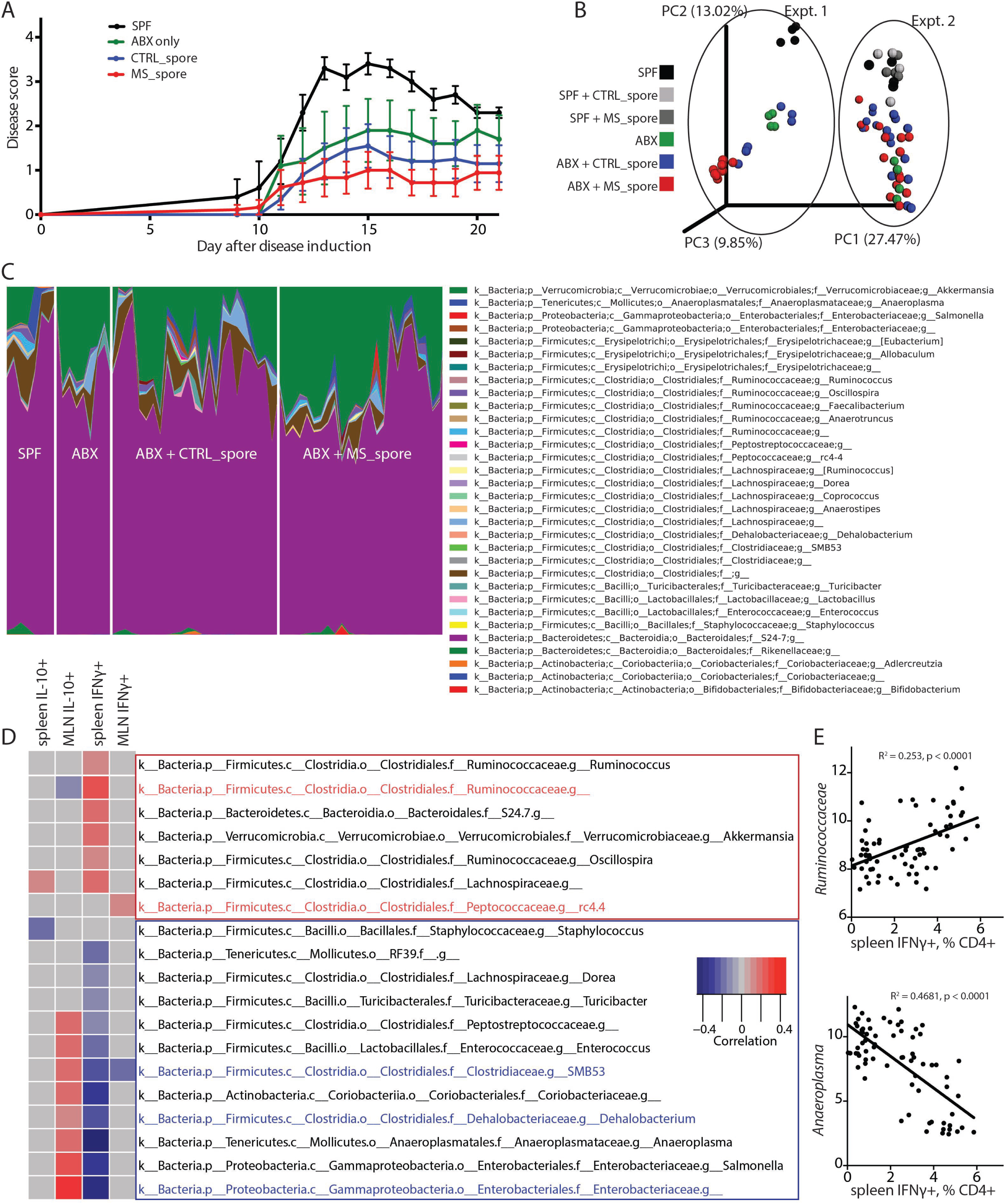
Spore-forming bacteria composition is correlated with T lymphocyte phenotypes *in vivo*. **A.** Clinical EAE scores of mice that after antibiotic treatment had been colonized with spore-forming bacteria from controls (CTRL_spore) or MS patients (MS_spore) for 2 weeks, or kept on antibiotics (ABX) or in SPF condition as controls, prior to induction of EAE at 9-10 weeks of age. N = 5-10 mice per group. **B, C.** Principal coordinate plot of beta diversity (PCoA; unweighted Unifrac) (**B**) and genus-level taxonomical distribution (**C**) of mouse fecal microbiota at 2 weeks of colonization with spore-forming bacteria, 2 separate experiments. **D.** Bacterial genera whose abundance is correlated with changes in immune cell differentiation in gnotobiotic mice are shown. The linear correlation between relative abundances of bacterial genera and the percentage of IL-10+ regulatory and IFNγ+ Th1 out of CD4+ Th lymphocytes from both spleens and mesenteric lymph nodes (MLN) of mice colonized with spore-forming bacteria are depicted as a heatmap. Same samples as in **B-C.** Only the genera that show significant linear correlation with immune parameters (p>0.05 after Benjamini-Hochberg adjustment for multiple comparisons) are included in the heat map. Red rectangle, putative pro-inflammatory subset. Blue rectangle, putative anti-inflammatory subset. Red font, taxa significantly increased in mice colonized with spore-forming bacteria from MS patients compared to controls. Blue font, taxa significantly reduced in mice colonized with spore-forming bacteria from MS patients compared to controls. **E.** Examples of positive and negative correlation between bacteria and Th lymphocyte differentiation from **D**.

We next analyzed whether spore-forming bacteria regulated T lymphocyte responses *in vivo*. To this end we colonized antibiotic-treated mice with spore-forming bacteria from 3 controls and 3 MS patients and analyzed the resulting changes in bacterial composition and T lymphocyte differentiation. Principal coordinate analysis (PCoA) of the beta diversity of gut microbiota separated SPF mice from antibiotic-treated and colonized (i.e. gnotobiotic) mice. While no major shifts in community composition based on disease state of the donor were observed (Fig. 3B), multiple microbial taxa were differentially abundant (Fig. 3C, Suppl. Tables 2, 3), including an increase in *Akkermansia* (3 OTUs corresponding to *A. muciniphila;* Suppl. Table 3) in mice colonized with spore-forming bacteria from MS patients.

Further investigation identified individual taxa that were classified as either pro-inflammatory or anti-inflammatory-based on the correlation between their relative abundance in mouse stool samples and their ability to alter differentiation of IFNγ+ Th1 or IL-10+ regulatory lymphocytes from either spleen or mesenteric lymph nodes (MLN) *in vitro* (Fig. 3D, 3E). The pro-inflammatory category (Fig. 3D, red rectangle) included taxa significantly increased in mice colonized with spore-forming bacteria from MS patients compared to controls (highlighted in red), while the anti-inflammatory category (mostly evident in splenocytes; blue rectangle) contained taxa significantly reduced in mice colonized with spore-forming bacteria from MS patients (highlighted in blue).

The increase in *Akkermansia muciniphila*, a non-spore-forming bacteria, in gnotobiotic mice colonized with spore-forming bacteria from MS patients led to the hypothesis that spore-forming bacteria may regulate *Akkermansia* levels. The correlation between spore-forming community composition and relative abundance of *Akkermansia* is shown in Fig. 4A. The increase in *Akkermansia* was present not only in the mice colonized with spore-forming bacteria from MS donors, but also in MS donors themselves (p = 1.5E^-09^, negative binomial Wald test) (Fig. 4B). Of interest, we and others (9, 10) recently reported the increased abundance of *Akkermansia* in untreated MS patients and identified this bacterium as sufficient for driving T lymphocyte differentiation into the pro-inflammatory IFNγ+ Th1 phenotype in-vitro (10). Consistent with this result, we also observed a significant positive correlation between the relative abundance of *Akkermansia* and IFNγ+ Th1 lymphocyte differentiation (Fig. 4C) in gnotobiotic mice. While other taxa also correlated with *Akkermansia* levels and T lymphocyte differentiation (Fig. 4D) our data suggest that the observed immunological effects may be at least partially mediated by *Akkermansia*.

**Figure 4.**
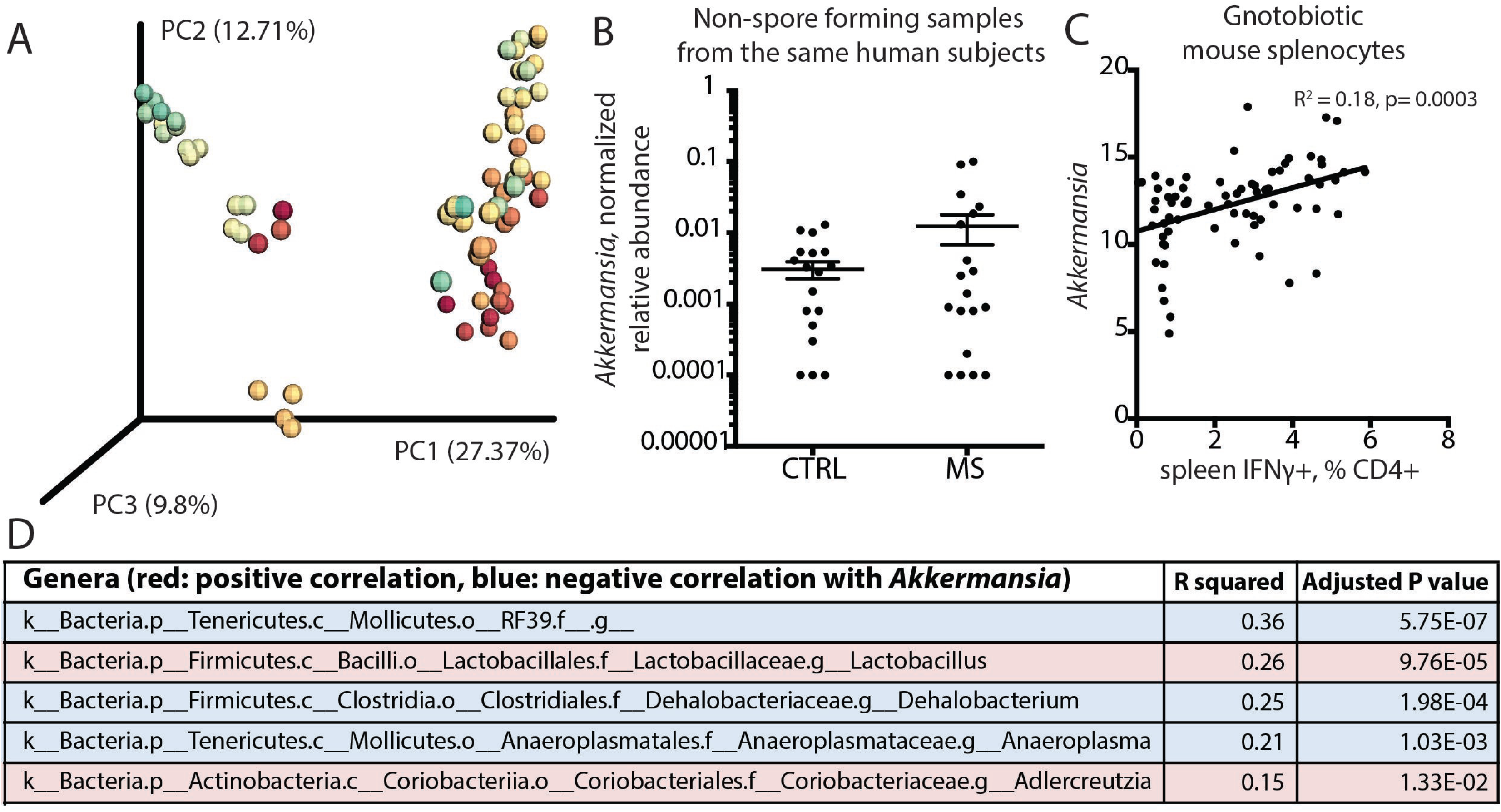
Increased *Akkermansia* is linked with MS-associated changes in spore-forming bacteria and pro-inflammatory T lymphocytes. **A.** Principal coordinate plot of beta diversity (PCoA; unweighted Unifrac) of mouse fecal microbiota excluding *Akkermansia* at 2 weeks of colonization with spore-forming bacteria, 2 separate experiments, colored by *Akkermansia* presence (red to green: low to high). p < 0.001, significant contribution of *Akkermansia* presence to determining distance variation (adonis method for continuous variables). **B.** Relative abundance of *Akkermansia* in controls and MS patients used for isolation of spore-forming bacteria. p = 1.5E-09, negative binomial Wald test, Benjamini-Hochberg correction for multiple comparisons (across all 144 species detected in the dataset). **C.** Linear correlation of relative abundance of *Akkermansia* with IFNγ+ Th1 lymphocyte differentiation in spleens of mice colonized with spore-forming bacteria. R^2 = 0.18, p= 0.0003. **D**. Bacterial genera significantly correlated with *Akkermansia in vivo*.

## Discussion

The spore-forming fraction of gut bacteria has been associated with immunoregulatory properties (4). Here we examined the structural composition and immunological effects of the spore-forming fraction of gut microbiota from subjects with MS compared to controls. MS-associated differences in bacterial community composition were correlated with impaired anti-inflammatory functions, as evidenced by a reduction in their ability to drive T lymphocyte differentiation into IL-10+ Tregs *in vitro*. Meanwhile, colonizing antibiotic-treated mice with spore-forming bacteria allowed us to identify specific taxa correlated with T lymphocyte differentiation into IFNγ+ and IL-10+ subtypes *in vivo.*

Our results contribute to the evidence supporting the immunoregulatory functions of spore-forming bacteria and show that these functions may be compromised in the context of autoimmunity. Previous studies on spore-forming bacteria had been conducted by isolating this fraction from a single healthy donor (4, 22). This approach allowed focusing on donor-specific bacterial strains, but provided limited information about the “baseline” composition and variability of this bacterial community in healthy humans. Here we used multiple healthy control donors to establish the baseline community composition of spore-forming bacteria, and compared these healthy profiles with those from patients with MS.

Our data corroborate previous findings that spore-forming bacteria, almost exclusively belonging to the phylum *Firmicutes,* and classes *Clostridia* and *Bacilli,* induce anti-inflammatory T lymphocytes *in vitro* and protect from autoimmune inflammation *in vivo* (4, 5). We also show that the taxonomical distribution and immunoregulatory functions of spore-forming bacteria are altered in MS patients. While were able to show that these differences have functional consequences in-vitro, they were not sufficient to alter the course of EAE using antibiotic treated mice. One possible explanation for this counterintuitive finding is that since our mice were treated with antibiotics, they were not completely germ-free prior to colonization. As a consequence, unexpected interactions among antibiotic resistant communities and the spore-forming fraction may have influenced the course of EAE. We recognize that using GF mice for these experiments could address some of these concerns. However, raising GF animals is still a highly specialized enterprise only available at select institutions. Further studies of gene expression and metabolic output of spore-forming bacteria may provide therapeutic targets for regulating T lymphocyte responses to reduce autoimmune inflammation.

The mechanisms by which spore-forming bacteria regulate host T lymphocyte differentiation remain to be discovered. Interestingly, an overlapping subset of bacterial taxa has recently been shown to inhibit host proteases, including cathepsins (23), which mediate adaptive immune responses by increasing Th17 (24) and limiting Treg differentiation (25). Although future studies are needed to establish this firmly, it is possible that spore-forming bacteria from controls, but not MS patients are able to stimulate Treg responses via cathepsin inhibition.

Furthermore, healthy human spore-forming bacteria produce short chain fatty acids (SCFAs), including butyrate and acetate (26), which have been observed to stimulate Treg and inhibit Th1 differentiation *in vitro* and *in vivo* (27), Mizuno 2017). Either pure butyrate or butyrate-producing spore-forming bacteria from healthy humans have been shown to be sufficient Treg induction (28) in mice. Thus, human T lymphocyte differentiation into Tregs may be driven by a yet-undiscovered SCFA-synthesizing subset of spore-forming bacteria that is present in controls and absent in MS patients.

*Akkermansia muciniphila* has previously been reported to be increased in MS patients compared to controls (9, 10, 29) and to have pro-inflammatory functions *in vitro* (10). In addition, *Akkermansia* has been shown to be resistant to broad-spectrum antibiotics (30), which in part may explain its persistence in mice colonized with spore-forming bacteria. The fact that high levels of *Akkermansia* were only seen in mice colonized with MS chloroform-resistant bacteria suggests that its population is normally regulated by commensals that are depleted in MS thus enabling *Akkermansia* overgrowth.

Our finding that *Clostridium perfringens* is more abundant in the spore-forming bacterial fraction of MS patients is consistent with the association of *C. perfringens* with neuromyelitis optica (NMO), another demyelinating autoimmune disease (31-33). Putative mechanisms of *C. perfringens*-mediated autoimmunity include molecular mimicry between *C. perfringens* peptide and a self-antigen in the human host (Varrin-Doyer 2012), and toxin-mediated increase in neuronal damage (32, 34).

Due to the high variability of spore-forming bacteria across donors, mouse colonization with samples from additional donor pairs would be required to assess whether MS-associated reduction in regulatory T lymphocyte differentiation *in vitro* can be reliably reproduced *in vivo.* However, a major advantage of gnotobiotic mouse models is the ability to assess the association between immune responses and microbial abundance within experimental communities. The identification of additional taxa capable of inducing clear differentiation paths in immune cells will further contribute to our understanding their role in immune regulation. For example, our findings corroborate the anti-inflammatory functions of relatively unknown bacterial genera such as *Anaeroplasma* and *Dehalobacterium* in mouse models of inflammation (35, 36).

In conclusion, we have investigated the immune functions of the spore-forming fraction of human gut microbiota in health and disease, using MS as a model of autoimmune inflammation. We identified novel bacterial taxa associated with MS as well as with T lymphocyte differentiation into both pro-inflammatory and regulatory phenotypes. Further studies of spore-forming bacteria and other experimentally defined bacterial populations may reveal specific immunoregulatory mechanisms in MS and other diseases that may be targeted by therapeutic interventions.

## Acknowledgements

We thank all subjects who participated in this study. Funding was provided by a grant (CA_1072-A-7) from the National MS Society. This study was also supported by a generous gift from the Valhalla Charitable Foundation. S.E.B is the Heidrich Family and Friends Endowed Chair in Neurology.

## Materials and Methods

### Isolation of spore-forming bacteria from human fecal samples

Fecal samples were collected from 25 adult patients with RRMS that had not received disease-modifying or steroid treatment for at least 3 months prior to the time of collection and 24 subjects without MS or any other autoimmune disorder (controls) at the University of California, San Francisco (UCSF) (Table 1). The inclusion criteria specified no use of antibiotics or oncologic therapeutics in 3 months prior to the study. All individuals signed a written informed consent in accordance with the sampling procedure approved by the local Institutional Review Board. Samples were stored in collection vials (Fisher #NC9779954) at -80° C until bacterial isolation.

**Table 1.**
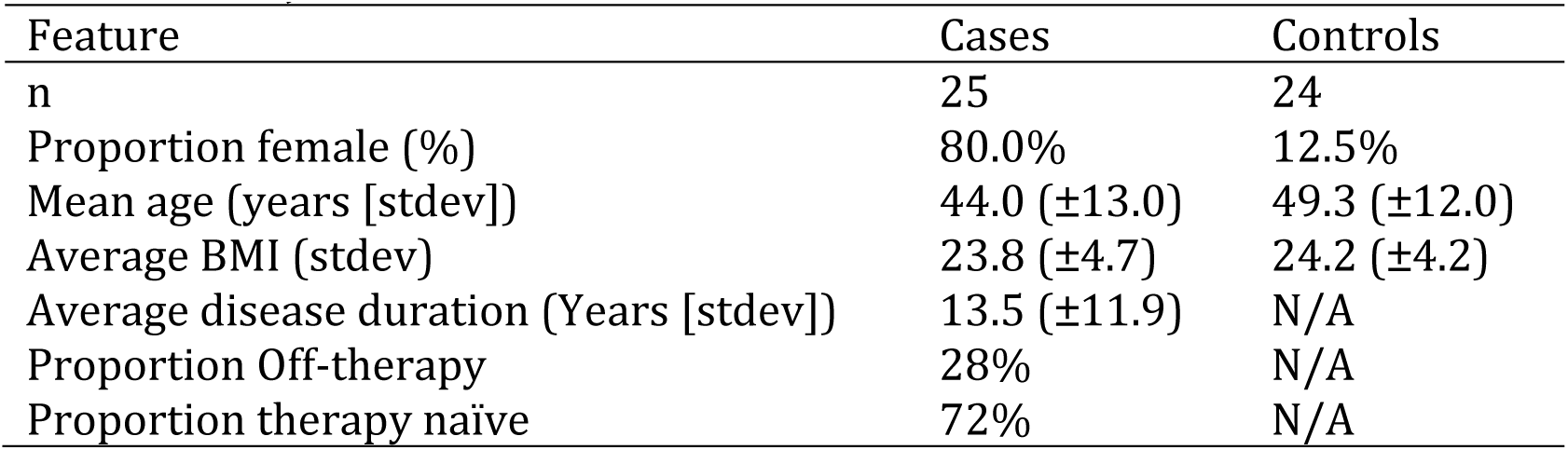
Subject characteristics

Spore-forming bacteria were isolated based on their resistance to chloroform as described previously (Atarashi 2013). Briefly, total bacteria were isolated from stool samples by suspending ~0.5mg stool sample in 1.5ml PBS, passing it three times through a 70µm cell strainer and washing twice with 1.5ml PBS by spinning at 8000rpm. The resulting suspension was diluted in 5ml PBS, mixed with chloroform to the final concentration of 3%, and incubated on a shaker for 1h at room temperature. After incubation, chloroform was removed from the solution by bubbling nitrogen (N2) gas for 30min. Chloroform-treated bacteria were then cultured on OxyPRAS Brucella Blood Agar plates (Oxyrase #P-BRU-BA) for 96 hours followed by Brucella Broth (Anaerobe Systems #AS-105) for 48 hours, and isolated for sequencing, *in vitro* experiments and *in vivo* experiments.

### 16S rRNA amplicon sequencing and computational analysis

DNA was extracted from mouse fecal or human chloroform-resistant bacterial culture samples using MoBio Power Fecal DNA extraction kit (MoBio #12830) according to manufacturer’s instructions. For each sample, PCR targeting the V4 region of the prokaryotic 16S rRNA gene was completed in triplicate using the 515/806 primer pair, and amplicons were sequenced on NextSeq at the Microbiome Profiling Services core facility at UCSF using the sequencing primers and procedures described in the Earth Microbiome Project standard protocol (11).

Analysis was performed using QIIME v1.9 as described (12). Essentially, amplicon sequences were quality-filtered and grouped to “species-level” OTUs via SortMeRNA method (13), using Greengenes v.13.8 97% dataset for closed reference. Sequences that did not match reference sequences in the Greengenes database were dropped from analysis. Taxonomy was assigned to the retained OTUs based on the Greengenes reference sequence, and the Greengenes tree was used for all downstream phylogenetic community comparisons. OTUs were filtered to retain only OTUs present in at least 5% of samples and covering at least 100 total reads. After filtering, samples were rarefied to 10000 sequences per sample. Alpha diversity was calculated using the Chao1 method (14). For analysis of beta diversity, pairwise distance matrices were generated using the phylogenetic metric unweighted UniFrac (15) and used for principal coordinate analysis (PCoA). For comparison of individual taxa, samples were not rarefied. Instead, OTU abundances were normalized using variance-stabilizing transformation and taxa distributions were compared using Wald negative binomial test from R software package DESeq2 as described previously (16, 17) with Benjamini-Hochberg correction for multiple comparisons. Linear correlations between bacterial taxa and lymphocyte proportions were computed after variance-stabilizing transformation of bacterial abundances (16).

### Mouse colonization with microbiota

Female littermates 5 week old C57BL/6J mice (JAX #000664) were treated with 1% solution of Amphotericin B in drinking water for 3 days, followed by 2 weeks of a solution composed of 1% Amphotericin B, 1mg/ml ampicillin, 1mg/ml neomycin, 1mg/ml metronidazole and 0.5mg/ml vancomycin in drinking water. After 2 weeks, the drinking solution was replaced by sterile water and mice were gavaged with specific bacteria of interest at 2*10^8 CFU in 100ul per mouse every 2 days for 2 weeks (7 total gavages). Bacterial colonization was either followed by the induction of EAE or immunophenotyping of mesenteric and cervical lymph nodes.

To induce EAE, mice were immunized in both flanks with 0.1ml _MOG35-55_ emulsion (1.5 mg/ml) mixed with Complete Freud’s Adjuvant and killed *Mycobacterium tuberculosis* H37Ra (2mg/ml), followed by two 0.1ml intraperitoneal injections of pertussis toxin (2µg/ml) immediately and at 48h after MOG/CFA injections. Mice were scored daily in a blinded fashion for motor deficits as follows: 0, no deficit; 1, limp tail only; 2, limp tail and hind limb weakness; 3, complete hind limb paralysis; 4, complete hind limb paralysis and at least partial forelimb paralysis; 5, moribund.

At the time of euthanasia, mouse mesenteric lymph nodes, and spleens were dissected and processed by grinding tissues through a 70µm cell strainer. Entire mesenteric and cervical lymph nodes and 10^7 splenocytes per mouse were stimulated for 4-5 hours with 20ng/ml PMA and 1µg/ml ionomycin in presence of protein transport inhibitor (GolgiPlug, BD #51-2301KZ) and used immediately for immunophenotyping, while the remaining splenocytes were stored for *in vitro* bacterial stimulations.

### Bacterial stimulation of human immune cells

Human peripheral blood mononuclear cells were isolated from healthy volunteers and stored at - 80° C in cryovials at 10^7 cells/ml concentration in FBS containing 10% DMSO. Before plating, cells were washed in PBS twice, re-counted, and plated at 10^6 cells/ml concentration in RPMI media supplemented with 10% FBS and 1% penicillin/streptomycin/glutamine. Cells were stimulated for 3 days as described previously (18) with anti-human CD3 (BD #555336, 0.3 µg/ml), anti-human CD28 (BD #555725, 2 µg/ml) and recombinant human TGF-β1 (R&D #240B002, 2.5ng/ml).

Bacteria isolated from human chloroform-resistant cultures were resuspended in PBS supplemented with protease inhibitor (Roche #4693159001) and phosphatase inhibitor (Roche #4906845001), heat-inactivated at 65° C for 1h and sonicated for 10min as described previously (19). Protein concentration in the resulting suspension was measured using the Pierce BCA protein assay kit (Thermo Scientific #23227). Bacterial extracts were added to PBMCs at 1µg/ml 1h after plating as described previously (20). PBS with the same protease inhibitor and phosphatase inhibitor was added as the no-bacteria control. Each human *in vitro* experiment contained at least 6 independent donor bacterial samples and was repeated at least twice.

### Immunostaining, flow cytometry and FACS of human immune cells

Human PBMCs were immunostained using standard protocols. Live/dead cell gating was achieved using Live/Dead Fixable Aqua kit (ThermoFisher #L34957). FoxP3/transcription factor staining buffer set (eBioscience #00-5523-00) was used for staining of intracellular and intranuclear cytokines. The following antibodies were used for human PBMC staining: antiCD3-PE.Cy7 (BD #563423), anti-CD4-PerCP.Cy5.5 (BioLegend #300530), anti-CD25-APC (BD #555434), anti-FoxP3-AlexaFluor488 (BD #560047) and anti-IL-10-PE (eBioscience #12-7108).

Flow cytometry was performed on BD Fortessa cell analyzer and analyzed using FlowJo software (TreeStar). Cells were gated to identify the lymphocyte population based on forward and side scatter, followed by gating for single color and live cell populations. Fluorescence minus one (FMO) was used for gating. Unstained, single color and fluorescence-minus-one controls were used to identify stained populations. For T lymphocyte suppression assay, control CD4+ CD25+ lymphocytes were sorted from PBMC cultures incubated with extracts from unrelated control or MS spore-forming bacteria in Treg-differentiating conditions on Aria III cell sorter (BD Biosciences) and cultured with CD4+ CD25- from the same donor pre-loaded with a CFSE cell division tracker kit. Statistical significance of expression changes in markers of T lymphocyte differentiation and proliferation was determined using two-tailed Student’s *t* test to compare samples from different donors and two-tailed repeated measures *t* test to compare samples from the same donor. GraphPad Prism 6 software was used to analyze and plot the data. *P* < 0.05 was considered statistically significant.

### Data availability

All 16S amplicon sequencing data presented in this article are available from the corresponding author upon request.

## Supplementary Material

**Supplementary Table 1. Spore-forming OTUs that were significantly different between MS patients and controls.** Negative binomial Wald test with Benjamini-Hochberg correction for multiple comparisons.

**Supplementary Table 2. Genera that were significantly different between antibiotic-treated mice colonized with spore-forming bacteria from MS patients and controls.** Negative binomial Wald test with Benjamini-Hochberg correction for multiple comparisons.

**Supplementary Table 3. OTUs that were significantly different between antibiotic-treated mice colonized with spore-forming bacteria from MS patients and controls.** Negative binomial Wald test with Benjamini-Hochberg correction for multiple comparisons.

**Supplementary Figure 1. Relative abundance of *Clostridium perfringens* OTUs in spore-forming bacteria of MS patients and controls.** N=30 patients, 24 controls. X axis, OTU IDs taken from GreenGenes 13.8 database. Y axis, relative abundances after rarefaction to 10,000 reads/sample. Last two columns (highlighted on graph) represent the sum of all individual OTUs.

